# Anxiety attenuates learning advantages conferred by statistical stability and induces loss of volatility-attuning in brain activity

**DOI:** 10.1101/2021.11.21.469465

**Authors:** Elise G. Rowe, Clare D. Harris, Ilvana Dzafic, Marta I. Garrido

## Abstract

Anxiety can alter an individual’s perception of their external sensory environment. Previous studies suggest that anxiety can increase the magnitude of neural responses to unexpected (or surprising) stimuli. Additionally, surprise responses are reported to be boosted during stable compared to volatile environments. Few studies, however, have examined how learning is impacted by *both* threat and volatility. To investigate these effects, we used threat-of-shock to transiently increase subjective anxiety in healthy adults during an auditory oddball task, in which the regularity could be stable or volatile, while undergoing functional Magnetic Resonance Imaging (fMRI) scanning. We then used Bayesian Model Selection (BMS) mapping to pinpoint the brain areas where different models of anxiety displayed the highest evidence. Behaviourally, we found that threat-of-shock eliminated the accuracy advantage conferred by environmental stability over volatility in the task at hand. Neurally, we found that threat-of-shock led to both attenuation and loss of volatility-attuning of neural activity evoked by surprising sounds across most subcortical and limbic brain regions including the thalamus, basal ganglia, claustrum, insula, anterior cingulate, hippocampal gyrus and also the superior temporal gyrus. Conversely, within two small clusters in the left medial frontal gyrus and extrastriate area, threat-of-shock boosted the neural activity (relative to the safe and volatile condition) to the levels observed during the safe and stable condition, while also inducing a loss of volatility-attuning. Taken together, our findings suggest that threat eliminates the learning advantage conferred by statistical stability compared to volatility. Thus, we propose that anxiety disrupts behavioural adaptation to environmental statistics, and that multiple subcortical and limbic regions are implicated in this process.

## INTRODUCTION

Anxiety affects both clinical (Hao et al., 2020) and non-clinical populations (Cao et al., 2020; Roy et al., 2020; Taylor et al., 2020). Non-clinical, transient anxiety has overlapping features with pathological anxiety, suggesting that studying the former can aid our understanding of the latter (Grillon, 2008; Robinson et al., 2013). A common way to study anxious states in healthy individuals are threat-of-shock paradigms, such as those used in several key neuroimaging studies of anxiety (Klumpers et al., 2017; Torrisi et al., 2018). Using such paradigms, Cornwell and colleagues (2007) found that healthy individuals under transient threat-of-shock show threat-induced increased neural responses to unexpected (or deviant) stimuli, comparable to people with post-traumatic stress disorder (Morgan and Grillon, 1999). We refer to this phenomenon of threat-induced increased neuronal responses as *hypersensitivity*. Interestingly, a later study by the same group (Cornwall et al., 2017) found that this hypersensitivity to unexpected stimuli could be reversed using anxiolytic medication (known to alleviate anxiety) administered during a combined threat-of-shock and auditory oddball paradigm.

Oddball paradigms use changes in the auditory (or visual) sensory statistics to evoke prediction error responses that are driven by larger neural responses to surprising (compared to frequent or unsurprising) events (Nätäänen, 1992). While a small number of studies have shown that these neural responses to surprising events are boosted during anxious states (Cornwell et al., 2007; 2017), it remains unclear how the stability or volatility of the environmental statistics affect such responses. Previous work has shown that neurotypical individuals increase their learning rates in volatile compared to stable environments (Behrens et al., 2007), and that anxiety detrimentally affects how quickly adults learn about new environmental contingencies following a surprising change in sensory statistics (Browning et al., 2015). We refer to this as threat-induced *loss of volatility-attuning*. In comparison, less is known about how anxiety affects an individual’s *accuracy* in their judgements relating to changing environmental statistics and which brain regions are engaged in these processes. As such, we investigated the degree to which induced anxiety, within a combined threat-of-shock and oddball paradigm, affects both the learning accuracy and brain responses to surprising stimuli in stable and volatile environments during threatening (threat-of-shock) and safe conditions.

Motivated by the above notions of threat-induced *loss of volatility-attuning* (Browning et al., 2015) and *hypersensitivity* (Cornwell et al., 2017), we designed a study to empirically test these hypotheses and which were operationalised as families of models (see Figure 2 for further information). We defined ‘volatility-attuning’ as the contextual modulation of surprising responses dependent on the stability of the environment (this neural definition is subtly different to that of Browning et al., 2015). Accordingly, an individual with intact volatility-attuning would display greater responses to deviant stimuli during the stable versus volatile conditions even under threat. Thus, a loss of volatility-attuning induced by threat would be reflected in models incorporating equal neural responses in volatile and stable contexts. Conversely, hypersensitivity during threat (Cornwell et al., 2017) was reflected in models that postulated greater activity for surprise responses under threat than safety, with or without a loss of volatility-attuning. Alternative models hypothesised *attenuation* with threat, meaning larger responses during safe conditions. These hypotheses were based on previous findings of reduced neural activity in response to unexpected sounds for post-traumatic stress disorder (PTSD) patients compared to controls (McFarlane et al., 1993; Menning et al., 2008).

**Figure 1:**
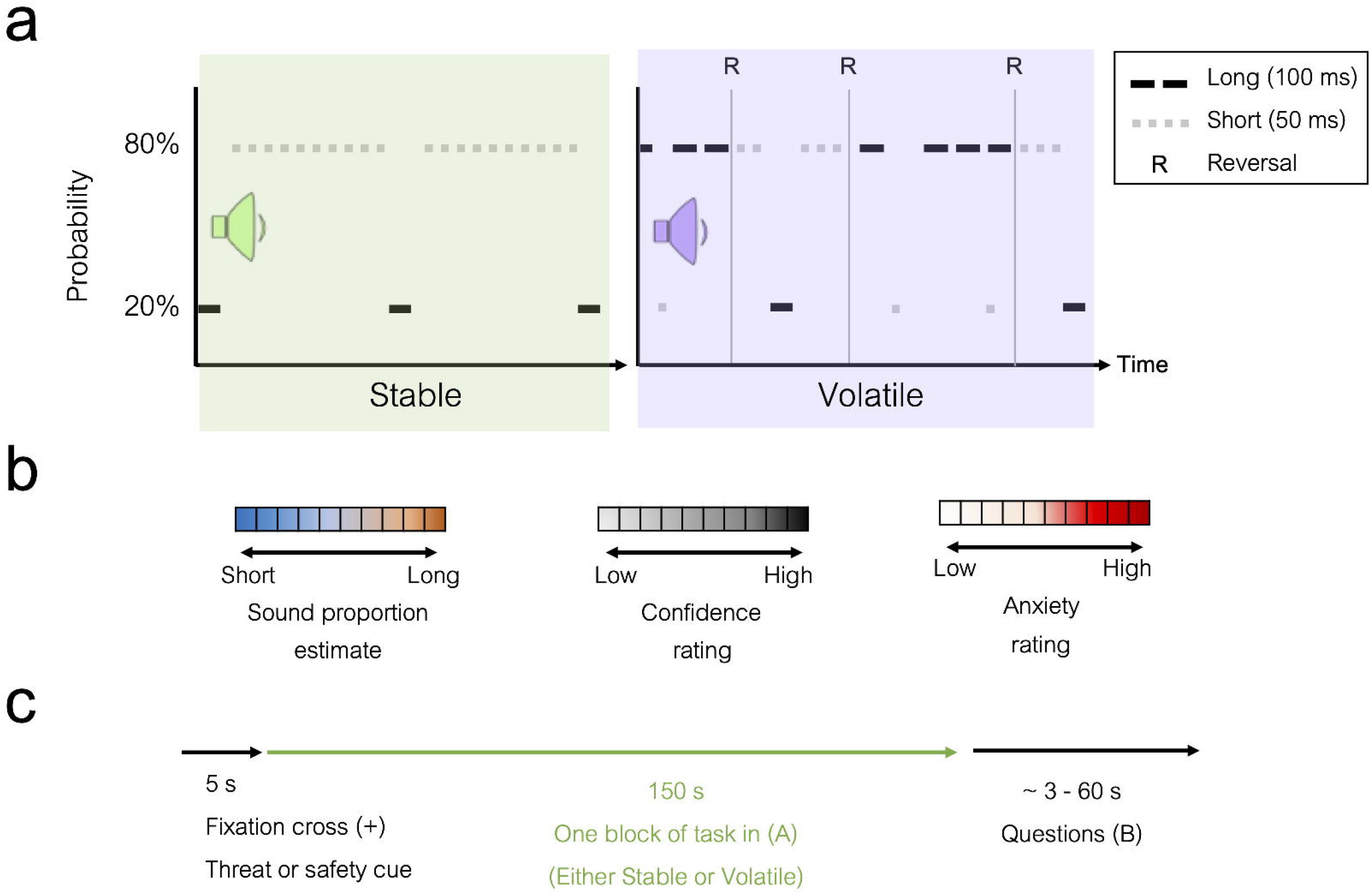
Threat-of-Shock combined with Auditory Volatility Oddball (TSAVO) Experimental Paradigm. **(a)** Example of one statistically stable (green) followed by one volatile (purple) block. Long dashed black lines denote long sounds while short dashed grey lines denote short sounds. In the volatile block, the letter “R” denotes “reversal” moments, when probability rules were reversed (e.g., from 80/20% to 20/80% for the long and short tones). This occurred three times during each volatile block. Each block was followed by three questions (see b). **(b)** Participants answered three questions after each block: (1) sound proportion estimate of the frequencies of the sounds (with ‘more short sounds’ displayed to the left hand side of the scale and ‘more long sounds’ on the right)), (2) their confidence rating (from low to high), and (3) a rating of their anxiety level (from low to high). **(c)** Overview of the components of each block. Each block began with either a threatening or safe cue, and ended with the same three questions. During threatening blocks, a single shock occurred at a variable, unpredictable length of time within 100 seconds from the end of the block.

**Figure 2:**
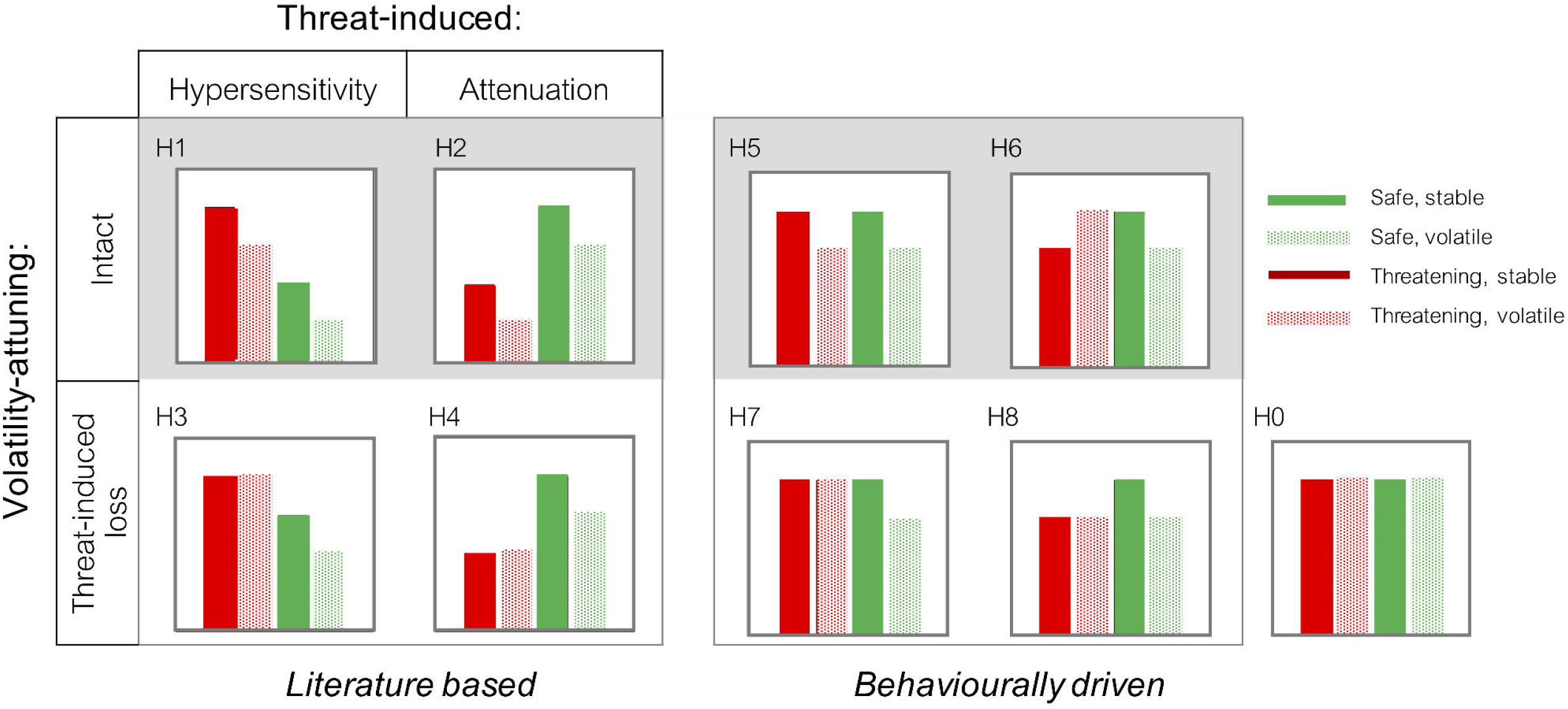
The nine hypotheses regarding the neural responses to surprising (deviant) stimuli. For our Bayesian Model Selection analysis, we formulated two groups of hypotheses that were either based on the literature (H1-H4) or behaviourally driven (H5-H8). All hypotheses could be grouped according to whether volatility-attuning (i.e., higher responses during stable compared to volatile conditions) remained intact under threatening conditions (H1, H3, H5, H6) or was lost (H2, H4, H6, H8). The literature-based models could be further divided according to whether neural responses to surprise were consistently highest in threatening conditions (H1, H3) or safe conditions (H2, H4). The behaviourally-driven models were all based on the behavioural statistical learning error (SLE) data. Finally, a null hypothesis (also referred to in the text as Hypothesis 0, H0) was included to reduce the Type I error rate. Please see the text for further explanations of each model.

To test these hypotheses we used Bayesian Model Selection (BMS) mapping for functional Magnetic Resonance Imaging (fMRI, Rosa et al., 2010). The BMS methodology is ideally suited to our purposes as it allows for the simultaneous comparison of any number of models at each and every voxel throughout the brain. BMS is unique as it is not limited to simply using contrasts to determine where an experimental effect is the greatest (as in traditional methods). Instead, one can determine where each of the hypothesised models is most likely. The results from our BMS analysis revealed where in the brain our different hypotheses regarding the anxiety-induced neural responses were most probable given our experimental data.

## MATERIALS AND METHODS

### Participants

Our final sample included 38 healthy participants (50% male, 50% female), ranging from 18 to 31 years of age (M = 21 years, SD = 2.89). All participants were right-hand dominant, fluent English speakers, with normal or corrected-to-normal vision. Participant exclusion criteria included the current use of psychotherapy or of psychotropic medication/s, excessive alcohol consumption (>2/day average), frequent tobacco use (>6/day), any contraindications to magnetic resonance imaging (MRI) scans, any history of mental or neurological disease, or an unwillingness to experience discomfort. The final sample participants’ demographics and relevant questionnaire scores are shown in Table 1. All participants gave both written and verbal informed consent to the study and were compensated for their time at a rate of $20 AUD per hour. The University of Queensland Institutional Human Research Ethics committee approved the study.

**Table 1:** Participant Demographics and Inventory Scores.

| Characteristic | N | Mean (SD) |
| --- | --- | --- |
| Sex | Male: 19; Female: 19 |  |
| Age (years) | 38 | 21 (2.89) |
| Trait Anxiety Score | 37* | 38 (8.12) |
| Beck’s Anxiety Inventory Score | 38 | 8 (8.24) |
| Beck’s Depression Inventory Score | 37* | 7 (7.19) |
| Intolerance of Uncertainty | 38 | 25 (7.95) |
| Statistical Learning Errors | 38 | 14 (5.18) |

\* Note that one participant declined to complete each of these questionnaires.

### Participant exclusion

From an original sample size of 52 individuals, we removed 14 participants. Five were excluded due to exclusionary medical conditions that were not disclosed during participant screening, six due to equipment failure or data loss, and three due to their behavioural results strongly suggesting they were not engaging in the task. We also note that following the experiment, 18 participants reported that they heard distorted sounds in some blocks. However, after determining there were no significant differences in the behavioural data with those who did not report the distorted sounds, we did not exclude any participants on this basis.

### Task Description and Procedure

We adapted the reversal oddball paradigm developed by Dzafic and colleagues (2020) from electroencephalography (EEG) to functional Magnetic Resonance Imaging (fMRI), and combined this with a modified version of a threat-of-shock procedure (Cornwell et al., 2017). This “Threat-of-Shock with Auditory Volatility Oddball” (TSAVO) task is depicted in Figure 1. Prior to the task, all participants completed an MRI metal check, changed into medical scrubs, and completed two to four practice blocks (depending on if they needed or requested more practice) of the task. All participants demonstrated understanding of the task prior to participating, and were provided opportunities for questions both before and after the task. The total time of the task was approximately 30-36 minutes, including short breaks.

The auditory stimuli consisted of pure tones delivered binaurally via earphones. Each tone was 1000 Hz in frequency and was either “short” (50 ms) or “long” (100 ms), with an interstimulus interval (ISI) of 750 ms. Although this ISI was fixed, there was a variable length of time for questions/responses after each block, which produced a “jitter” between blocks.

There were 12 blocks per experiment, with 200 tones delivered per block (each block lasting 150 seconds). The tones were played in a pseudorandom order, with the “standard” being common (occurring with 80% probability), and the “deviant” being uncommon, or unpredictable. Each deviant always had at least one standard tone played after it. In half the blocks, the frequency of standard and deviant tones were kept constant throughout the block (80/20%): these were called *stable* blocks (Figure 1a, left). During *volatile* blocks, the standard and deviant tone probabilities were reversed at three unpredictable times per block, as depicted in Figure 1a (right). We counterbalanced the presentation order of the four types of blocks (stable safe, volatile safe, stable threatening, and volatile threatening) across participants.

While Cornwell et al. (2007, 2017) only administered one shock despite multiple shock warnings, in this experiment participants were administered a single shock once during five of the six “shock-warning blocks”. Each “shock block” had a differently-timed shock, delivered at an unpredictable time *t* ≤100 seconds from the end of the block. We randomised the shock delivery time because temporally unpredictable administration of aversive stimuli seems to increase the anxiety induced by said stimuli (e.g., Grillon et al., 2004; Somerville et al., 2012; Simmons et al. 2008). Thus, the aim was for participants to be relatively sure they were likely to receive a shock, but to be unable to predict *when* they would receive it.

Prior to the experiment, each participant’s anterior wrist surface was cleaned and two electrodes were placed approximately 2 cm from the wrist flexion crease. Next, participants were administered a work-up shock procedure during which between one and five sample shocks were delivered to the wrist via a constant current stimulator. Shocks were delivered via a Biopac machine which functioned in association with *Acqknowledge Data Acquisition* software and *MATLAB*. Participants were asked to choose a shock intensity that was physically uncomfortable but not painful. Their chosen level of shock, a moderately uncomfortable physical intensity, was kept constant throughout the experiment. The shock level ranged from 0.001 to 50 V for this participant group, with most participants selecting a voltage between 0.01 to 0.02 V. Before each block, a message appeared on the screen saying either, “PREPARE FOR SHOCK AT ANY TIME” (“threat cue”) or “YOU ARE SAFE. NO SHOCK THIS ROUND” (“safe cue”).

### Behavioural measure of Statistical Learning Errors (SLEs)

After each block, participants estimated the percentage of the short and long sounds, as well as their confidence and anxiety levels (both rated from 0 to 100), using visual rating scales (Figure 1b). For each block, we calculated the behavioural *Statistical Learning Errors* (SLEs), which we defined as the absolute difference between a participant’s estimate of the percentage of a given type of sound, and the true percentage of those sounds. For example, if the percentage of short sounds was 20% and a participant responded with 40%, then their SLE was 20%. We also calculated each participant’s average stable-over-volatile learning advantage in the safe or threatening blocks, which we defined as the difference in SLEs (in percent) between stable and volatile conditions. The confidence and anxiety ratings were taken to be the location along the visual scale where the participant clicked, converted to a percentage of the total scale length (with 0% on the left/lower end of the scale, and 100% on the right).

### Data acquisition and pre-processing

Structural and functional MRI images were acquired on a 3-T Siemens Magnetom Prism scanner using a 32-channel head coil. Prior to the task, structural images were acquired with an MP2-RAGE sequence with a repetition time (TR) of 4000 ms, echo time of (TE) of 2.91 ms, resolution (voxel size) of 1 mm^3^, Field of View (FoV) of 256 mm, 176 slices per slab, inversion times (TI and T2) of 900 ms and 2220 ms respectively, and flip angles of 6 degrees and 7 degrees respectively. During the task, functional T2*-weighted images were acquired using a multiband, echo-planar sequence, across the whole brain (TR: 785 ms, TE: 30.0 ms, resolution: 2 mm^3^ isotropic, slices: 60, FoV: 208 mm, flip angle: 52 degrees). Following the completion of the task, there was also the collection of diffusion imaging data using a neurite orientation dispersion and density imaging (NODDI) technique (Zhang et al., 2012); analyses of those data will be reported elsewhere.

Standard pre-processing of the fMRI images was completed using the Statistical Parametric Mapping 12 (SPM12; http://www.fil.ion.ucl.ac.uk/spm/) software package for MATLAB (The MathWorks, Inc.; http://www.mathworks.com). The pre-processing steps included realignment on the functional images; co-registration of the functional and structural images; segmentation of the structural image, with heavy regularisation (0.1) recommended for MP2-RAGE sequence; normalisation of the resliced images into a standardised, stereotaxic space (according to the Montreal Neurological Institute, MNI, template); and smoothing of normalised images with an 8mm full-width-at-half-maximum isotropic Gaussian kernel.

### Behavioural analyses

Behavioural analyses were performed using Two-Way Analyses of Variance (ANOVAs) for the main effects and interactions of volatility and threat on anxiety ratings and the SLE data. These were followed up with post-hoc tests, with significance values adjusted using Bonferroni correction for multiple comparisons. We also examined the *difference* in the SLE learning advantage conferred by stable over volatile environments by subtracting each participant’s average SLE in the volatile condition from the safe condition, separately within the safe or threatening conditions. Significance was examined using a paired-sample t-test.

### Bayesian Model Selection

A Bayesian approach allowed us to compare multiple non-nested hypotheses operationalised as different models of the neural activity under the different volatility and threat conditions. We specified and compared nine different hypotheses (named H0 to H8, and shown in Figure 2) using random effects (RFX) Bayesian Model Selection (BMS) as described by Rosa et al. (2010) and the SPM12 manual (Ashburner et al., 2016). For a full description of the BMS method we employed, including the use of parametric modulators (i.e., regressor weights) to specify different hypotheses about the relative amplitude of neural responses, please see Rosa et al. (2010) and Harris et al. (2018). Briefly, for each model, parametric modulators were assigned to each experimental condition such that they represented the hypothesised relative amplitude of the neural response to a deviant event. For example, if we hypothesised that, for condition A, the neural activity levels were 4 times greater in magnitude than in condition B, then we would assign a weighting of “4” for condition A and a “1” for condition B. In this example, the parametric modulators would simply consist of a vector of “4”s (one for each trial) in condition A, and a vector of “1”s for the trials in condition B.

#### BMS model definitions

Our intention was to describe plausible, easily interpretable hypotheses for the relative deviant neural responses under different combinations of threatening, safe, volatile, and stable conditions. It is important to note that, when using model comparison, the most probable model is conditioned upon *those contemplated in the model space*. It is often intractable to test for all possible models. Fortunately, however, the literature and our behavioural findings aided in narrowing the (technically unlimited) range of possible models down to eight simple alternative hypotheses and a null. We drew on findings in the literature to inform the models representing Hypotheses 1 to 4 (H1-H4), and we drew inspiration from our behavioural data, in combination with a previous study (Dzafic et al., 2020), to construct the models representing Hypotheses 5 to 8 (H5-H8). Finally, we constructed a null hypothesis, H0, to directly assess the probability that there were no differences between the four conditions. For each of these hypotheses, we included every model that reflected all possible combinations of integer parametric modulators (between 1 and 4) representing the hypothesized ranking of neural activity amongst the conditions (see below and Supplementary Material for further details). We limited the parametric modulators to integers between 1 and 4 for simplicity. We describe how we specified these models below, where we refer to the experimental conditions of safe stable (SS), safe volatile (SV), threatening stable (TS) and threatening volatile (TV).

We designed H1 to H4 based on the previous literature. H1 was informed by the aforementioned findings of Cornwell et al. (2007, 2017) and Dzafic et al., (2020), with greater neural activity under threat than in safety (hypersensitivity) and during stable compared to volatile conditions (intact volatility-attuning). H1 was specified using parametric modulator weights of 4, 3, 2 and 1 for TS, TV, SS and SV conditions, respectively. As there were no other possible combinations of integers from 1 to 4 to represent the rankings of neural activity between the experimental conditions, H1 included only one model alternative. H3 was a variation of H1 reflecting hypersensitivity to threat, but with loss of volatility-attuning during threat, inspired by the behavioural findings of Browning et al. (2015). H3 had the parametric modulator weights of 4, 4, 3 and 2 for TS, TV, SS and SV conditions, respectively. In addition to this ‘base’ version of H3, there were 3 other sets between 1 and 4 that could represent the same ranking of neural activity amongst the conditions (i.e., [4,4,2,1], [3,3,2,1] and [4,4,3,1]), giving a total of four H3 model variations.

H2 and H4 were based on previous findings of reduced neural activity (or *attenuation*) in response to unexpected sounds in post-traumatic stress disorder (PTSD) patients compared to controls (McFarlane et al., 1993; Menning et al., 2008). These models were derived by modifying H1 and H3, respectively, such that neural responses were greater in *safety* than in threat, again combined with either retention (H2) or loss (H4) of volatility-attuning during threat. For H2, the regressor weights were 2, 1, 4 and 3 for the TS, TV, SS and SV conditions, respectively. There were no further possible sets of integers from 1 to 4 that could specify H2. For H4, the weights for the ‘base’ model were 2, 2, 4, 3 for the TS, TV, SS and SV conditions, respectively. In addition, there were 3 variations of regressor weights specifying the same rank order of neural activity (i.e., [1, 1, 4, 3], [1, 1, 3, 2] and [1, 1, 4, 2]), giving a total of four variations of H4. Overall, for H1-H4, there were a total of 10 models.

In contrast to these hypotheses, H5 to H8 were based on the patterns observed in the behavioural SLE data. We created these models to test whether there were regions whose neural deviant responses varied in a way that was proportional to behavioural accuracy (that is, inversely proportional to SLEs). Our results (see Results for further details) showed that during safe volatile conditions the SLEs were significantly higher than during the safe stable conditions; meaning participants made more SLEs during the safe volatile blocks (compared to the safe stable blocks). As such, we hypothesised that the deviant neural responses would be greater (suggesting greater prediction errors) in the safe stable blocks than the safe volatile blocks. These models were inspired by previous findings that surprise responses tend to be larger for people with higher behavioural accuracy (Dzafic et al., 2020). Given this, we designed four hypotheses (H5-H8) that matched the observed significant difference whereby the response in SS conditions was always larger than the response in SV conditions.

In H5 the conditions were represented as SS = TS > SV = TV (using parametric modulators of 4, 3, 4, 3 for the TS, TV, SS and SV conditions, respectively), denoting intact volatility-attuning under threat. In H6 the conditions were represented as SS = TV > SV = TS (using parametric modulators of 3, 4, 4, 3 for the TS, TV, SS and SV conditions, respectively), also denoting preserved volatility-attuning. In H7 the conditions were represented as SS = TS = TV > SV (using parametric modulators of 4, 4, 4, 3 for the TS, TV, SS and SV conditions, respectively). H7 also reflected a loss of volatility-attuning with threat and hypersensitivity in volatile environments (with respect to the safe and volatile condition). Finally, in H8 the conditions were represented as SS > SV = TS = TV (using parametric modulators of 3, 3, 4, 3 for the TS, TV, SS and SV conditions, respectively). H8 also reflected a loss of volatility-attunning during threat, as well as attenuation of the neural responses in safe environments (with respect to the safe and stable condition). As with the models of H1-H4, we included all possible regressor weight combinations representing the same rank ordering. For each of the models H5-H8 there were 5 possible combinations of integers from 1 to 4 (inclusive), giving a total of 20 model variations within this behaviourally-inspired model set. Finally, H0, the null, had the same parametric modulator weight (4) for all four conditions. Hence, our nine hypotheses were operationalised as 31 different models when considering all of the above mentioned variations of parametric modulators.

In our application of BMS, an estimate of the log of the model evidence (log-model evidence) was calculated for each model at every voxel, for each participant. These were all compared using RFX BMS which allowed us to determine the (potentially different) ‘winning model’ (i.e., the model with the highest log model evidence given the data) at each voxel within the brain. After this, the exceedance probability maps were displayed with user-specified probability thresholds of 0.85. This enabled the visualisation of the different regions of the brain where different models most likely explained the fMRI data; that is, with >85% probability.

#### Bayesian Model Maps

All displayed results have a minimum cluster size of 5 voxels. The EPMs show the exceedance probability which is an estimate of the relative probabilities that any given model explains the data (in the listed voxels) better than the other models considered. Note that we did not exclude white matter regions from our results. The physiological relevance of white matter signals in fMRI datasets is beyond the scope of this paper, but has been discussed elsewhere (Cheng et al., 2015; Gawryluk et al., 2014; Gore et al., 2019; Grajauskas et al., 2019). Also note that when displaying the EPMs, one can select any user-defined threshold. A frequentist may wish to display EPMs where the exceedance probability is at least 95%, but if you do so, note that the interpretation of these results is different (and arguably much simpler): clusters displaying above the 95% EPM threshold have at least 95% exceedance probability compared to all the other models considered, given the data collected. In our results, we choose to present any EPMs above 85%. We present our BMS results using the BrainNet Viewer software package for Matlab (Xia et al., 2013) and label the anatomic regions using the Anatomy Toolbox add-on for SPM 12 (Eickhoff et al., 2005; Zilles & Amunts, 2010; Amunts et al., 2007) and the Multi-Image Analysis Graphical User Interface (GUI) (Mango) software.

## RESULTS

### Behavioural findings

#### Anxiety scores increased following threat-of-shock blocks

We examined differences in participants’ average anxiety ratings (from 0 to 100, recorded here as %) following safe blocks compared to threatening blocks, and in stable blocks compared to volatile blocks. We found a significant main effect of threat, F(1,37) = 16.32, *p* < 8.55×10^−5^, with higher reported anxiety ratings following threatening blocks (M = 42.84%, SE = 3.36) compared to safe blocks (29.05%, SE 3.41), with a mean rating difference of 13.78%. There was no significant main effect of volatility, F(1,45) = 0.01, *p* = 0.943, nor was there an interaction effect of threat and volatility on anxiety ratings, F(1,45) = 0.06, *p* = 0.805.

#### Statistical learning errors are lower during stable compared to volatile blocks, but only under safe *conditions*

We estimated the SLEs across the four experimental conditions. We used a two-way repeated-measures ANOVA to assess the significant differences between the groups and found there was a significant main effect of volatility, F(1,37) = 12.73, *p* < 4.85×10^−4^. Following this, we conducted post-hoc statistical tests, finding significantly higher SLEs in the volatile compared to stable conditions within safe blocks (*p* < 5.91×10^−5^ adjusted by Bonferroni correction for multiple tests; see Figure 3).

**Figure 3:**
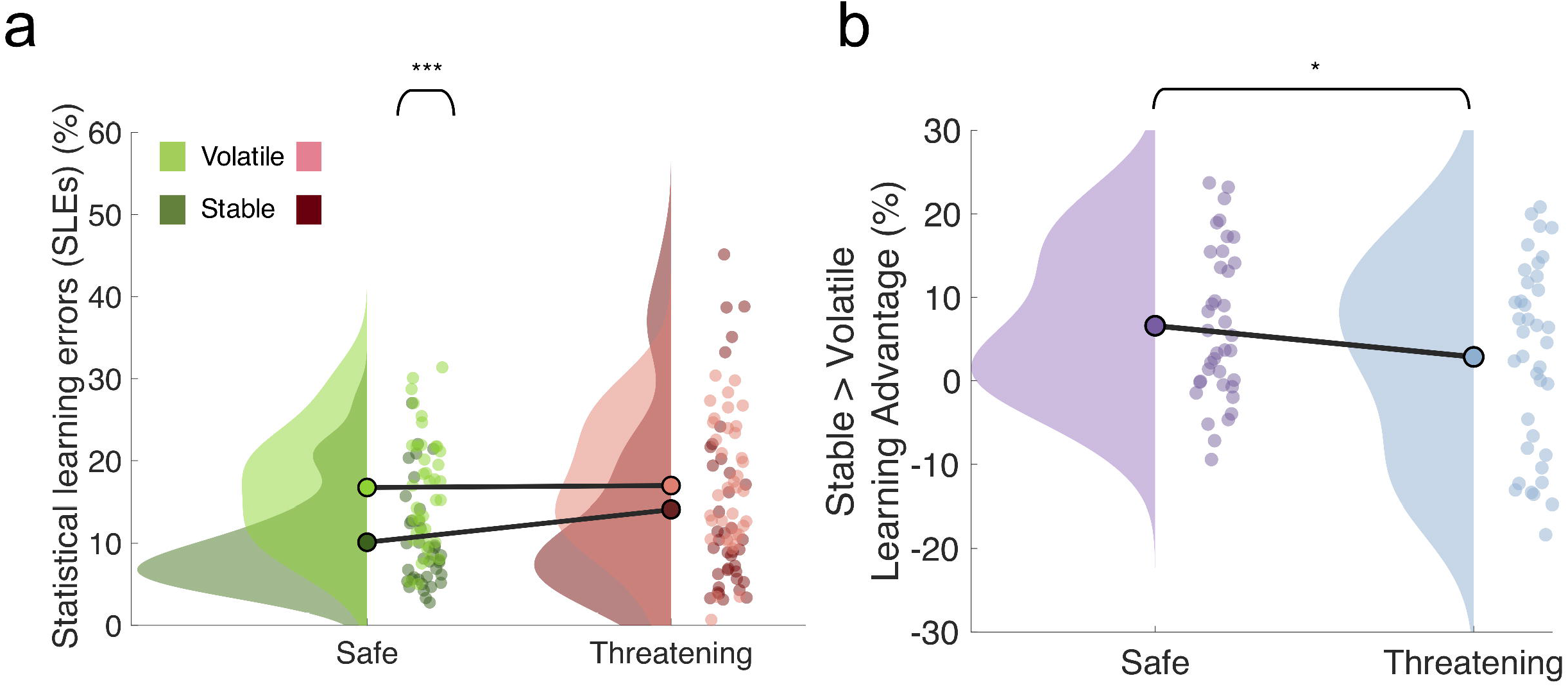
Statistical Learning Errors (SLEs) under different levels of volatility and threat. **(A)** Participants’ average SLEs are represented by density plots (each participant’s average is represented by a single dot in the scatter plots, and group mean SLEs are represented by connected, dark dots). In the safe and stable condition, participants made significantly fewer SLEs compared to the safe and volatile condition. **(B)** Participants’ average stable > volatile learning advantage is represented by a dot in the density plot. We observed a significant reduction in the learning advantage conferred by stable compared to volatile environments during threatening conditions. Note: The asterisks (***) and (*) are indicative of a significant difference of p < 0.0001 and p < 0.05, respectively.

#### Anxiety attenuates the learning advantages conferred by stability over volatility

Following the behavioural findings of other studies (Browning et al., 2015, Huang et al., 2017, Piray et al., 2019), we tested whether there was threat-induced mitigation of the learning advantages usually conferred by stable compared to volatile conditions. For this, we estimated the difference in the SLEs between the stable and volatile conditions within the safe or threatening blocks. The average SLE reduction conferred by statistical stability over volatility, was 6.60% in safety, compared to 2.87% in volatile conditions, showing that threat significantly attenuated the stability-over-volatility learning advantage (*p =* 0.033, t = 2.21, *df =* 37).

### Summary - Bayesian Model Selection

We used BMS to determine which of our hypotheses best explained the neuroimaging data at each voxel. Our Exceedance Probability Maps (EPMs, set at 85% probability, shown in Figure 4 and described below) helped to shed light on the neural computations most likely to occur in different brain regions under the different levels of threat and volatility. Overall, we found that H8 (SS > SV = TS = TV; specifically, M8e in the Supplementary Material) contained the winning model across the majority of the (mostly subcortical and limbic) brain regions. This model suggests that surprise responses are attenuated in stable environments during threat and that there is an overall loss of volatility-attuning (i.e., no difference in the responses evoked during stable or volatile environments under threat). The latter observation of a loss of volatility-attuning is consistent with other work (Browning et al., 2015; Huang et al., 2017; Piray et al., 2019). In addition, we found that H7 (SS = TS = TV > SV; specifically, M7a in Supplementary Material), which reflects greater neural activity under threatening and safe stable environments (compared to safe volatile environments), was most likely in two small clusters in V3/V4 and the posterior-medial frontal gyrus. This model is also consistent with a loss of volatility-attuning under threat. Finally, the null hypothesis (H0) was most likely in a number of cortical areas within the occipital, parietal, temporal and frontal areas. Below, we provide further details relating to the results from these BMS analyses.

**Figure 4:**
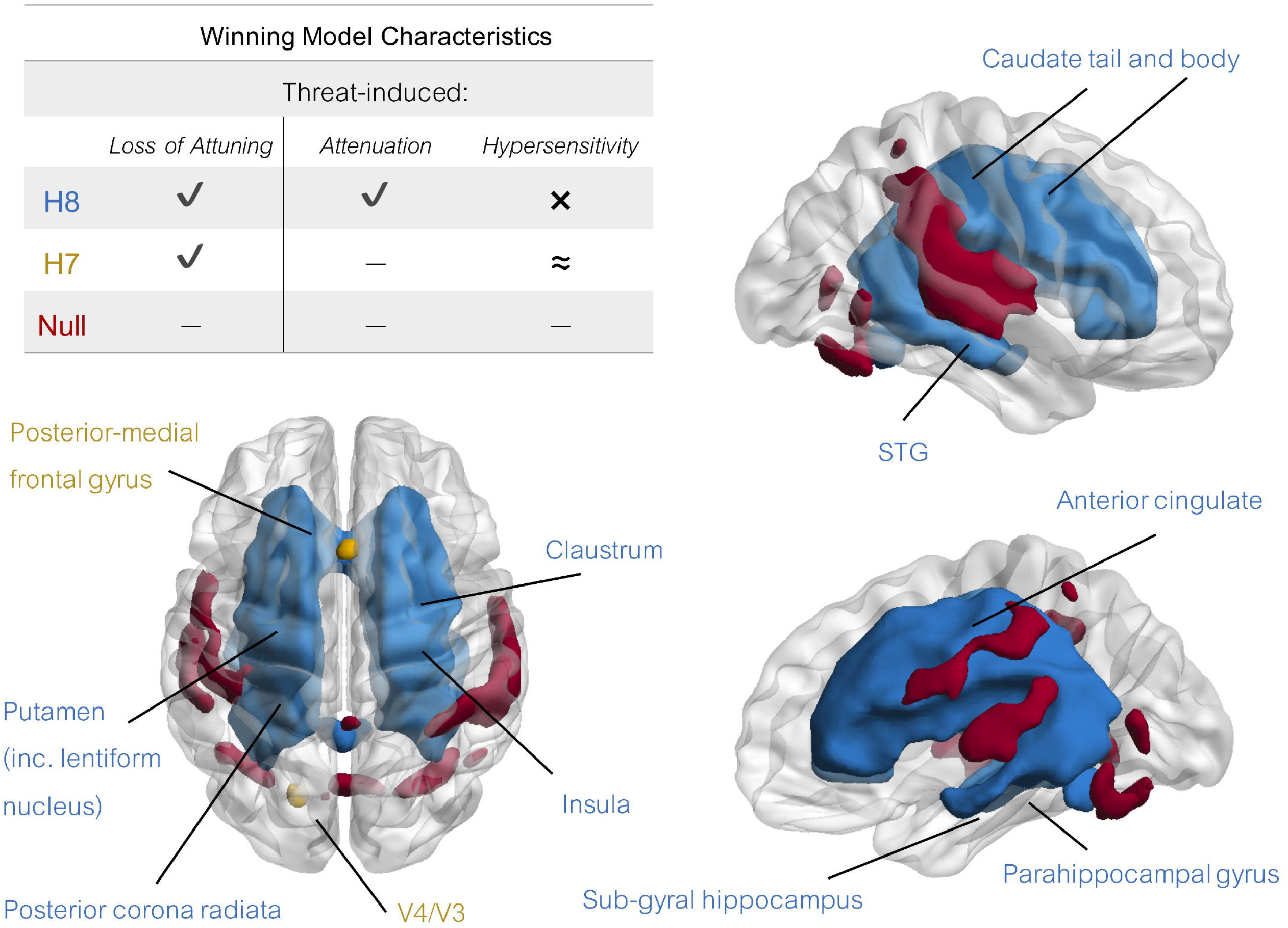
Exceedance probability maps (EPMs) thresholded at an exceedance probability of 0.85 for all models of each hypothesis. Selected clusters are labelled for Hypothesis 8, H8 (dark blue), Hypothesis 7, H7 (yellow) and Hypothesis 0, the ‘null’ (dark red). Hypothesis 1, (H1), 2 (H2), 3 (H3), 4 (H4), 5 (H5) and 6 (H6) did not have results above this threshold and are therefore not shown. Note: ‘Loss of Attuning’ refers to no difference between the stable or volatile deviant response during threatening blocks. ‘Attenuation’ refers to a reduction in the threat-based deviant responses (relative to the safe condition), while ‘Hypersensitivity’ refers to the opposite. Additionally, ‘**✔**’ = present, ‘x’ = absent and ‘≈’ = present but only within volatile conditions.

### Threat reduces neural activity and eliminates volatility-attuning in most subcortical brain regions

As discussed above and shown in Figure 4 and Table 2, H8 dominated most subcortical and limbic regions of the brain. This hypothesis stipulated that the greatest deviant neural activations are observed under stable and safe conditions, with threat *reducing neural activity* in stable environments and eliminating *volatility-attuning* (by eliminating differences in the neural activity otherwise seen between stable and volatile conditions). H8 (highlighted in dark blue in Figure 4) was far more likely than any other model, with over 99.99% exceedance probability over multiple regions, including bilateral caudate tail and body, parahippocampal gyrus, thalamus/putamen, lentiform nucleus of the putamen, anterior cingulate, the left STG and claustrum and right insula, sub-gyral hippocampus and posterior corona radiata.

**Table 2.** Hypothesis 8: Threat reduces neural activity and eliminates volatility-attuning.

| Brain region | Hem | BA | MNI coordinates |  |  | Cluster size | Peak |
| --- | --- | --- | --- | --- | --- | --- | --- |
|  |  |  | x | y | z |  | Exceedance |
|  |  |  |  |  |  |  | Probability (%) |
| Cingulum, STG, Caudate tail and body, parahippocampal gyrus, thalamus/putamen, lentiform nucleus of the putamen, claustrum, anterior cingulate, insula, sub-gyral hippocampus | L/R | NA | -16 | 12 | 34 | 14706 | 100.00 |
| Posterior corona radiata | R | NA | -6 | 44 | -10 | 151 | 99.00 |

*Note: This table contains the location for the peak of a very large EPM cluster with an $\geq 85\%$ probability threshold. Please note that the reported coordinate for the cingulum was simply the location of the peak exceedance probability, but the cluster included multiple other regions as listed here. L: left; R: right.*

#### Threat increases deviant neural responses in some frontal and occipital cortical regions

In two small clusters, we found that H7 displayed the greatest evidence. H7 proposed that neural activity would increase during *threatening* volatile blocks (see Figure 4 and Table 3) compared to *safe* volatile blocks, and included a loss of volatility-attuning. H7 had exceedance probabilities over 88% and 87% in small clusters in the left V4/V3 region (Brodmann Area/BA 19) and the left posterior-medial frontal gyrus (Brodmann Area/BA 46), respectively.

**Table 3.** Hypothesis 7: Threat increases neural activity and eliminates volatility-attuning.

| Brain region | Hem | BA | MNI coordinates |  |  | Cluster size | Peak |
| --- | --- | --- | --- | --- | --- | --- | --- |
|  |  |  | x | y | z |  | Exceedance Probability (%) |
| V4/V3 | L | 19 | -18 | -72 | -10 | 15 | 88.00 |
| Posterior-medial frontal gyrus | L | 46 | 2 | 20 | 44 | 10 | 87.00 |

*Note: This table lists Exceedance Probability Map (EPM) region clusters 5-or-more voxels in size with an $\geq 85\%$ probability threshold. Clusters in the same region are grouped together. Between clusters, the table order is determined by exceedance probability (and within clusters of the same probability, the order is determined by cluster size). L: left; R: right.*

### The Null Hypothesis

We included a null hypothesis in our set of models (Hypothesis 0, H0) as a way to protect against Type I errors when using RFX BMS to calculate exceedance probabilities (Moser et al., 2018; but see alternative methods, Rigoux et al., 2014; Correa et al., 2018). As listed in Table 4, H0 explained the data best in clusters in the bilateral central opercular and superior parietal lobe, the left occipito-temporal gyrus, V1 and frontal pole, and in the right lateral occipital cortex, V4, and temporal pole, suggesting that deviant responses in these regions were not sensitive to manipulations of threat or volatility.

**Table 4.** Model 0 (Null): Neural activity is unaffected by threat and volatility.

| Brain region | Hem | BA | MNI coordinates |  |  | Cluster size | Peak |
| --- | --- | --- | --- | --- | --- | --- | --- |
|  |  |  | x | y | z |  | Exceedance Probability (%) |
| Central opercular cortex | R | 42 | 58 | -6 | 8 | 1,473 | 100.00 |
|  | L |  | -58 | -18 | 10 | 936 | 100.00 |
| V4 | R | 19 | 28 | -72 | -14 | 250 | 100.00 |
| Lateral occipital cortex | R | 18 | 48 | -62 | -2 | 95 | 100.00 |
| Lateral occipito-temporal gyrus | L | 37 | -30 | -68 | -16 | 222 | 100.00 |
| Superior parietal lobe | R | 5 | 2 | -46 | 60 | 53 | 100.00 |
| V1 | L | 17 | -2 | -70 | 12 | 201 | 100.00 |
| Frontal pole | L | 10 | -44 | 36 | -6 | 6 | 94.00 |
| Superior parietal | L | 7 | -26 | -52 | 42 | 11 | 93.00 |
| Temporal pole | R | 38 | 52 | 8 | -4 | 9 | 90.00 |

*Note. This table lists EPM clusters 5-or-more voxels in size with an $\geq 85\%$ probability threshold.*
*L: left; R: right.*

## DISCUSSION

We investigated statistical learning in the context of both stable and volatile environments, during threatening and safe conditions (i.e., with and without the physical threat-of-shock, respectively). We found that statistical learning accuracy was higher during safe blocks when the environmental statistics were stable compared to volatile, and that the learning advantages conferred by stable environments were lost under threat. Next, to model the different surprise responses occurring simultaneously across the brain, we used BMS to compute the EPMs for nine hypotheses (operationalised as model families) that were based on either the previous literature or our behavioural findings (and including a null model). We found that H8 hypothesising threat-induced loss of volatility-attuning and attenuation of neural surprise responses (but only with respect to the stable conditions) best explained the data in the majority of (mostly subcortical and limbic) brain regions. In other words, during threatening blocks, most subcortical and limbic regions exhibited reduced deviant responses (i.e., attenuation) and no difference in the responses evoked during the stable or volatile conditions (i.e., loss of volatility-attuning). In two small clusters, that included V4/V3 and the posterior-medial frontal gyrus, we also observed threat-induced loss of volatility-attuning but here instead of threat-induced attenuation (as in H8) we observed an increased response during the threatening and volatile condition (H7). Finally, we found a number of cortical regions in which the deviant responses were unaffected by threat and volatility (i.e., where the null hypothesis won); these included clusters in the occipital, parietal, temporal and frontal regions.

### Subcortical and limbic regions display a loss of volatility-attuning and attenuation of surprise responses under threat

For most subcortical and limbic brain regions, responses to surprising stimuli were greatest during safe and stable blocks compared to the other three conditions (as shown in Figure 4, H8). According to H8, threat-induced attenuation occurred during stable but not volatile conditions. These effects were observed in many subcortical and limbic regions including the insula, claustrum, putamen, caudate, anterior cingulate, parahippocampal gyrus and the STG. Findings within areas such as the insula (Steuwe et al., 2014, 2015), and (dorsal) ACC (Klumpers et al., 2010) are in support of previous literature showing these areas are engaged in threat processing. More broadly, the role of these limbic structures in generating prediction error responses, particularly during states of anxiety, has been discussed in a number of recent reviews (Calhoon & Tye, 2015; Tucker & Luu, 2021). Here, the limbic system is referred to as the neural circuitry underlying anxiety and is credited for the assignment of emotional weighting to potential threats (Calhoon & Tye, 2015). Others have identified the limbic regions as a base for Bayesian prior formation and for organising predictive coding processes throughout the neocortex (Tucker & Luu, 2021). Our findings align with these recent proposals, and support the notion that the distributed sensory information that is processed through the limbic system leads to affectively charged prior expectancies which influence future updating of beliefs.

Interestingly, however, these threat-induced attenuation findings are in opposition to our original literature-based *hypersensitivity* hypotheses, H1 and H3, which were informed by multiple previous findings showing greater neural responses to surprising stimuli in anxiety. Notably, most of these studies were performed using EEG or magnetoencephalography (MEG; except for Chen et al., 2017, who used fMRI). Indeed, Morgan and Grillon (1999) found that people suffering from post-traumatic stress disorder had increased neural responses to surprise compared to controls (see also Ge et al., 2011; Bangel et al., 2017) and Chang et al. (2015) found the same in people with panic disorder. Similar findings were observed in people scoring high in trait anxiety (Chen et al., 2017) and among neurotypical individuals under threat-of-shock (Scaife et al., 2006; Cornwell et al., 2007, 2017). Despite being at odds with the hypersensitivity hypotheses, these findings are consistent with multiple other EEG studies, which show a pattern of *reduced* surprise responses in anxiety disorders, or ‘threat-induced attenuation’, including case-control studies in PTSD (McFarlane et al., 1993; Menning et al., 2008) and panic disorder (Tang et al., 2013; Rentzsch et al., 2019). We highlight a number of potential factors contributing to this below.

### Loss of volatility-attuning and hypersensitivity under threat in frontal and occipital brain regions

Despite finding no evidence for the literature-based models of generalised hypersensitivity-during-threat (i.e., H1 and H3), we did observe two small clusters in which threat-induced-hypersensitivity occured, but only under volatile environments (i.e., H7). These two clusters lay within the right posterior medial-frontal gyrus and the left V4/V3 area. The location of one of these clusters, the posterior medial-frontal gyrus, is consistent with previous hypersensitivity findings from Cornwell and colleagues (2007) auditory oddball paradigm with a threat-of-shock as well as with Klumpers et al.’s (2010) findings that this region responds more during threat. However, this hypersensitivity only occurred under volatile (not stable) environments. Moreover, H7 only explained the data in a relatively small number of voxels and we did not find any significant clusters for H1 and H3. Thus, our results do not support the traditional definition of hypersensitivity-during-threat. There are multiple potential factors that contributed to this. Firstly, our participants may have set lower shock levels than in other studies, since they knew their discomfort ratings would be used to set their shock level, unlike in Klumpers and colleagues’ (2010, 2017) experiments. Secondly, shocks were set at a maximally uncomfortable level without being painful, unlike in Browning and colleagues’ (2015) experiment. These factors might also explain the fact that our average anxiety ratings did not exceed the halfway point on the scale (in contrast to, e.g., Cornwell et al., 2017), even though they were greater following the threat-of-shock compared to safe blocks.

### Threat eliminates volatility-attuning both behaviourally and neurally

Our behavioural SLE results support our BMS findings by suggesting that threat reduces the learning advantages conferred by statistical stability compared to volatility (Figure 3b). Critically, this suggests that anxiety affects learning *accuracy* in similar ways to how it affects learning *rates*, as shown by previous studies (Browning et al., 2015, Huang et al., 2017, Piray et al, 2019). Our findings also support the effects of anxiety upon volatility-attuning, which has been explained in diverging ways by different research teams. On the one hand, Browning et al. (2015) showed that those high in trait anxiety under threat-of-shock display dysfunctional learning in volatile environments. The authors inferred that these individuals exhibited decision-making patterns that indicate difficulty in updating action-outcome plans based on the statistics of the current environment. Consistent with this idea, Raio et al. (2017) showed that anxiety specifically impaired reversal learning, although that study did not include multiple reversals and therefore did not introduce volatility in the way that our study did. Contrary to the conclusions above, other studies (Huang et al., 2017; Piray et al., 2019) have found that anxious people do not have impairments in learning in volatile environments *per se*, but instead display a failure in learning within stable environments. Our behavioural and neural (H8 and H7, see Figure 4) results speak strongly in support of the notion of a loss of volatility-attuning and the behavioural findings suggest that the threat-induced loss-of-volatility-attuning was caused by dysfunctional learning in stable environments (Huang et al., 2017; Piray et al., 2019).

### Neuroanatomical correlates of loss of volatility-attuning

The previously identified neuroanatomical correlates of the loss of volatility-attuning in anxiety are also consistent with the current study. For example, Piray et al. (2019) found that although the BOLD signal in the dorsal ACC (dACC) correlated with learning rate across all trials and participants, this was not the case for those high in social trait anxiety. Despite our focus on learning *accuracy* instead of learning *rates*, and that we used transient, mildly-elevated anxious states (compared to high trait social anxiety), we similarly found that in the ACC, threat led to loss of volatility-attuning (and attenuation) of neural surprise responses. However, it will be important for future work to replicate and clarify these results, especially considering that this is but one study, and considering the paucity of other studies investigating these topics to date.

As discussed above, similar learning effects have also been observed in highly trait anxious people under threat-of-painful-shock (Browning et al., 2015), and even in people deemed likely to have clinical anxiety compared to controls (Huang et al. 2017). Thus, similar patterns are emerging across the spectrum from transient anxiety in neurotypical people, through to trait and clinical anxiety. Taken together with the previous studies, these findings are consistent with the conceptualisation of anxiety as a dimensional construct (Fucci et al., 2019). We hope that future research can develop our understanding further, by examining learning and anxiety in larger samples of people, including the full spectrum from healthy to clinically anxious participants. Further investigations will also elucidate how neural responses and behavioural learning findings like ours, relate to underlying structural tracts (McFadyen et al., 2020) and functional circuits (Shackman & Fox, 2016; Fudge et al., 2017).

## CONCLUSION

In summary, our BMS findings show that increased subjective anxiety induced by threat-of-shock in healthy individuals leads to a loss of volatility-attuning across the majority of subcortical and limbic brain regions. In addition to a loss of attuning, we observed attenuation of neural activity under threat during stable conditions in the majority of subcortical and limbic regions of the brain. The loss of volatility-attuning and attenuation of neural responses to surprising events occurred within regions including the thalamus, basal ganglia, claustrum, insula, anterior cingulate, hippocampal gyrus, and the superior temporal gyrus. Conversely, threat-induced hypersensitivity (or increased activity) during volatile conditions was also observed in the left medial frontal gyrus and left extrastriate area, where the threat-of-shock boosted the neural activity to levels observed during the stable blocks. Overall, our results show that threat eliminates volatility-attuning by removing the learning advantages conferred by stable over volatile conditions – both on a behavioral level (in terms of learning accuracy) and on a neural computational level across most brain regions (as per H8 and H7). Our EPMs also shed light on the possible neuroanatomical underpinnings of similar behavioural findings in other studies. Finally, our results are also consistent with theories regarding the limbic system’s role in affective processing of sensory information, including the formation of prior expectations and the updating of beliefs. In the future, we hope that studies in clinical populations can build upon both our behavioural and neural findings, and in so doing, uncover potential targets for future interventions.

## Acknowledgments

We thank all participants for their time, the radiographers at the Centre for Advanced Imaging (Nicole Atcheson and Aiman Al-Najjar) for help with data collection, David Lloyd for technical assistance with programming the statistical learning task, and the staff providing MASSIVE high performance computing support (Goscinski et al., 2014).

