## Supplementary material for "Anxiety attenuates learning advantages conferred by statistical stability and induces loss of volatility-attuning in brain activity": Rowe, Harris, Dzafic & Garrido - SUPPLEMENTARY MATERIAL.pdf

**Authors:** Elise G. Rowe[a,b]*, Clare D. Harris[b,c]*, Ilvana Dzafic[a,b,c,d,e,f] and Marta I. Garrido[a.b,c,d]

**Affiliations**
[a]Melbourne School of Psychological Sciences, The University of Melbourne, Melbourne, Australia
[b]Australian Research Council Centre of Excellence for Integrative Brain Function, Australia
[c]Queensland Brain Institute, University of Queensland, Brisbane, Australia
[d]Centre for Advanced Imaging, University of Queensland, Brisbane, Australia
[e]Orygen, the National Centre of Excellence for Youth Mental Health, Melbourne, VIC, Australia
[f]Centre for Youth Mental Health, University of Melbourne, Australia
*These authors contributed equally to this work.

### Model Definitions

In our BMS analysis, we tested nine hypotheses (see Main Text Figure 2) regarding the relative magnitude of the neural responses to surprising stimuli (i.e., deviants) under the 2 levels of volatility (i.e., volatile and stable) and threat (i.e., threatening and safe). As discussed in the Methods section, for each hypothesis, we included every model that reflected all possible combinations of integer parametric modulators between 1 and 4. That is, we did not double-test models that were equivalent to each other (i.e., representing the same *relative* difference between the experimental conditions). For example, for four experimental conditions, the parametric modulators of [1,2,2,2] could also be represented as [2, 4, 4, 4]. Integers between 1 and 4 (inclusive) were chosen for simplicity, but it is noted that each hypothesis could be specified by a theoretically infinite number of models and the parametric modulator weights could be specified as anything, and need not be integers between 1 and 4. We included all combinations of parametric modulators because we wanted to rigorously test each model by comparing it against all possible integer parametric modulator combinations. Overall, we tested a total of 31 models across the 9 hypotheses (see Supplementary Table 1 below). Four of the nine hypotheses (Hypotheses 1 to 4, ‘H0’ to ‘H4’, comprising 10 sets of parametric modulators) were based on the literature, four (Hypotheses 5 to 8, ‘H5’ to ‘H8’, comprising 20 sets of parametric modulators) were based on the behavioural results of our task and a final (Hypothesis 0, ‘H0’) was the ‘null hypothesis’, representing no difference in the neural responses between the experimental conditions.

**Supplementary Table 1**

**Bayesian Model Selection (BMS) model space for the nine Hypotheses (H0 to H8) comprising 31 model variations.**

| | | SS | SV | TS | TV |
|---|---|---|---|---|---|
| H1 | a | 2 | 1 | 4 | 3 |
| H2 | a | 4 | 3 | 2 | 1 |
| H3 | a | 3 | 2 | 4 | 4 |
| | b | 2 | 1 | 4 | 4 |
| | c | 2 | 1 | 3 | 3 |
| | d | 3 | 1 | 4 | 4 |
| H4 | a | 4 | 3 | 2 | 2 |
| | b | 4 | 3 | 1 | 1 |
| | c | 3 | 2 | 1 | 1 |
| | d | 4 | 2 | 1 | 1 |
| H5 | a | 4 | 3 | 4 | 3 |
| | b | 3 | 2 | 3 | 2 |
| | c | 4 | 2 | 4 | 2 |
| | d | 4 | 1 | 4 | 1 |
| | e | 3 | 1 | 3 | 1 |
| H6 | a | 4 | 3 | 3 | 4 |
| | b | 3 | 2 | 2 | 3 |
| | c | 4 | 2 | 2 | 4 |
| | d | 4 | 1 | 1 | 4 |
| | e | 3 | 1 | 1 | 3 |
| H7 | a | 4 | 3 | 4 | 4 |
| | b | 4 | 2 | 4 | 4 |
| | c | 4 | 1 | 4 | 4 |
| | d | 3 | 2 | 3 | 3 |
| | e | 3 | 1 | 3 | 3 |
| H8 | a | 4 | 3 | 3 | 3 |
| | b | 4 | 2 | 2 | 2 |
| | c | 4 | 1 | 1 | 1 |
| | d | 3 | 2 | 2 | 2 |
| | e | 3 | 1 | 1 | 1 |
| H0 | a | 4 | 4 | 4 | 4 |

*Note. ‘SS’ = ‘Safe Stable’, ‘SV’ = ‘Safe Volatile’, ‘TS’ = ‘Threatening Stable’ and ‘TV’ = ‘Threatening Volatile’.*
